# Global transcriptional regulation of innate immunity in *C. elegans*

**DOI:** 10.1101/469817

**Authors:** Marissa Fletcher, Erik J. Tillman, Vincent L. Butty, Stuart S. Levine, Dennis H. Kim

## Abstract

The nematode *Caenorhabditis elegans* has emerged as a genetically tractable animal host in which to study evolutionarily conserved mechanisms of innate immune signaling. We previously showed that the PMK-1 p38 mitogen-activated protein kinase (MAPK) pathway regulates innate immunity of *C. elegans* through phosphorylation of the CREB/ATF bZIP transcription factor, ATF-7. Here, we have undertaken a genomic analysis of the transcriptional response of *C. elegans* to infection by *Pseudomonas aeruginosa*, combining genome-wide expression analysis by RNA-seq with ATF-7 chromatin immunoprecipitation followed by sequencing (ChIP-Seq). We observe that PMK-1-ATF-7 activity regulates a majority of all genes induced by pathogen infection, and observe ATF-7 occupancy in regulatory regions of pathogen-induced genes in a PMK-1-dependent manner. Moreover, functional analysis of a subset of these ATF-7-regulated pathogen-induced target genes supports a direct role for this transcriptional response in host defense. The genome-wide regulation through PMK-1– ATF-7 signaling reveals global control over the innate immune response to infection through a single transcriptional regulator in a simple animal host.

**Author Summary:** Innate immunity is the first line of defense against invading microbes across metazoans. *Caenorhabditis elegans* lacks adaptive immunity and is therefore particularly dependent on mounting an innate immune response against pathogens. A major component of this response is the conserved PMK-1/p38 MAPK signaling cascade, the activation of which results in phosphorylation of the bZIP transcription factor ATF-7. Signaling via PMK-1 and ATF-7 causes broad transcriptional changes including the induction of many genes that are predicted to have antimicrobial activity including C-type lectins and lysozymes. In this study, we show that ATF-7 directly regulates the majority of innate immune response genes upon pathogen infection of *C. elegans,* and demonstrate that many ATF-7 targets function to promote pathogen resistance.

## Introduction

Convergent genetic studies of host defense of *Drosophila melanogaster* and mammalian innate immune signaling revealed a commonality in signaling pathways of innate immunity that has helped motivate the study of pathogen resistance mechanisms in genetically tractable host organisms such as *Caenorhabditis elegans* [1]. The simple *C. elegans* host has enabled the genetic dissection of integrative stress physiology orchestrating host defense of *C. elegans*[2-5]. Genetic analysis of resistance of *C. elegans* to infection by pathogenic *Pseudomonas aeruginosa* has defined an essential role for a conserved p38 mitogen-activated protein kinase pathway that acts on a CREB/ATF family bZIP transcription factor, ATF-7, in immune responses [6,7]. A complementary approach to characterizing the host response has been organismal transcriptome-wide characterization of genes induced upon infection by a number of different bacterial pathogens [8-15]. Putative effector genes encoding lysozymes and C-type lectin domain (CTLD)-containing proteins have been identified that have also served as useful markers of immune induction. Here, we report the genome-level characterization of the *C. elegans* response to *P. aeruginosa* that is mediated by ATF-7 activity downstream of PMK-1 activation, combining RNA-seq analysis of pathogen-induced gene expression with ChIP-seq analysis of ATF-7 binding, which suggests global regulation of the immune response of *C. elegans* through a single MAPK- transcription factor pathway.

## Results and Discussion

We performed RNA-seq on wild-type (N2), *pmk-1* mutant, or *atf-7* mutant animals exposed to *E. coli* OP50 or *P. aeruginosa* PA14 to identify genes that are differentially regulated upon infection that also require PMK-1 or ATF-7 for induction (Figure 1A). We found that in wild-type animals, 890 genes were two-fold upregulated (adjusted p-value <0.05), and 803 genes were two-fold downregulated upon *P. aeruginosa* exposure, compared to animals exposed in parallel to *E. coli* (Figure 1B; Table S1). Many of these upregulated genes have been previously implicated in the *C. elegans* immune response and including genes encoding C-type lectin domain (CTLD)-containing genes and lysozymes, corroborating prior microarray-based gene expression studies (Figure 1C) [8-10,12-14]. In contrast, gene ontology analysis of genes that are decreased in expression upon *P. aeruginosa* exposure shows enrichment for genes associated with homeostasis with significant ontology terms consistent with growth, development and reproduction (Figure S1A). Of note, many of the genes upregulated in response to *P. aeruginosa* exposure exhibit relatively low expression when animals are propagated on *E. coli*, whereas genes that are decreased in expression upon *P. aeruginosa* exposure are expressed at a higher basal level during normal growth conditions on *E. coli* (Figure 1D, S1B). In parallel, we analyzed *P. aeruginosa*-mediated gene expression changes in *pmk-1* and *atf-7* mutants to identify the proportion of genes induced by *P. aeruginosa* exposure that required PMK-1 and/or ATF-7 for induction (Figure S2). We observed that 70% of genes significantly induced two-fold or greater by *P. aeruginosa* exposure were no longer fully induced upon loss of *pmk-1,* and that 53% of upregulated genes were no longer fully induced upon loss of *atf-7* (Figure 1E, Table S1). We also found that 41% of genes reduced two-fold or more by *P. aeruginosa* required PMK-1, and 50% required ATF-7 for reduction of expression (Figure S2B, Table S1). These data implicate a high degree of involvement of PMK-1-ATF-7 signaling in the majority of changes in gene expression induced in response to infection by *P. aeruginosa*.

**Figure 1:**
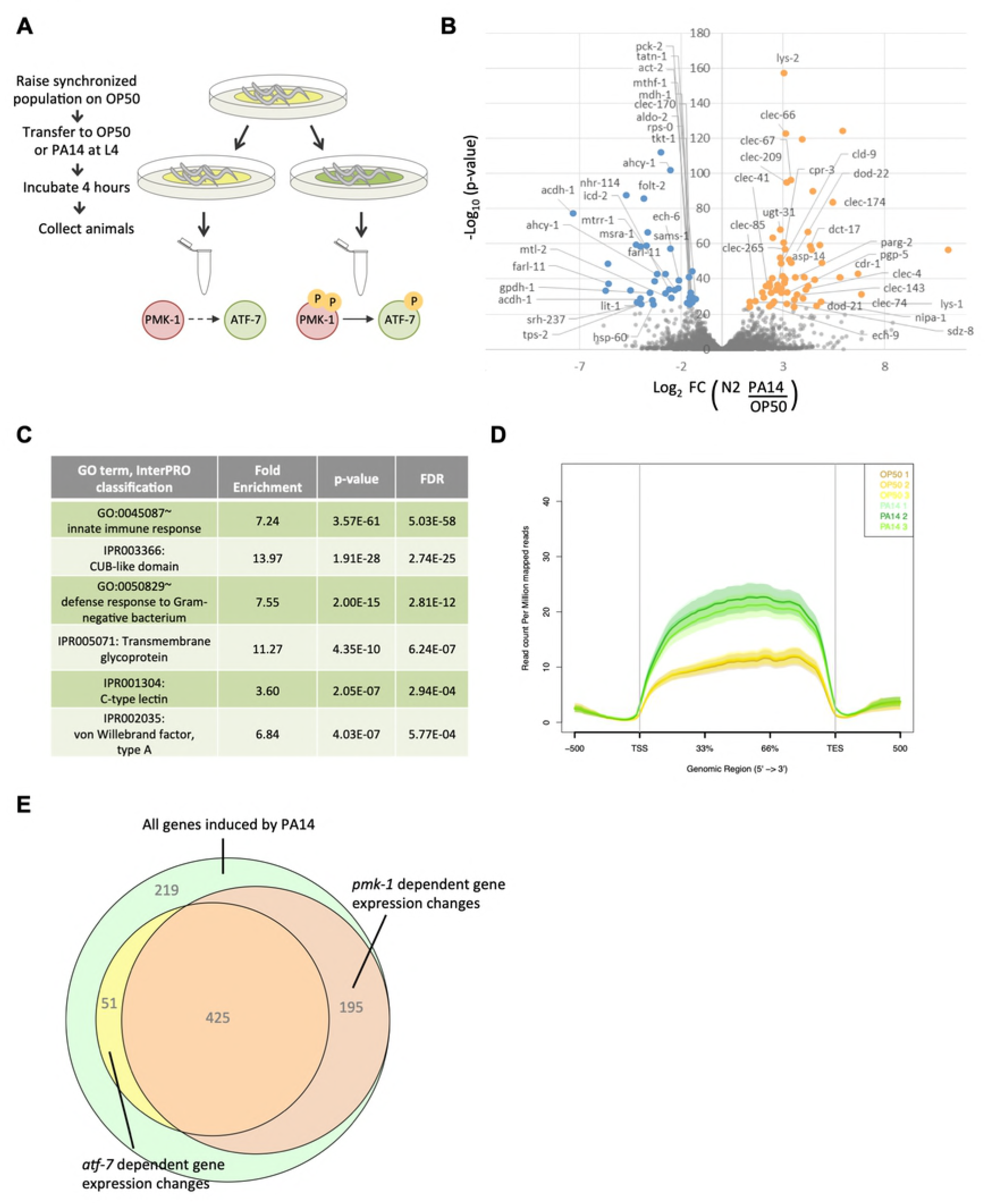
Exposure to *P. aeruginosa* prompts gene expression changes. **(A)** Schematic of experimental design. Yellow bacterial lawn indicates *E. coli* OP50, green bacterial lawn indicates *P. aeruginosa* PA14. PMK-1 and ATF-7 activation states are indicated below each condition. **(B)** Volcano plot of transcripts corresponding to differentially expressed protein-coding genes by exposure to *P. aeruginosa* PA14 versus *E. coli* OP50 in N2 animals. Orange and blue colored dots denote the top 100 outliers that are increased or decreased, respectively, and are annotated with their gene names. **(C)** Top GO terms and InterPRO classifications of transcripts that are significantly upregulated (adjusted p-value < 0.05) in N2 animals exposed to PA14 versus OP50. **(D)** Average expression (RPM) across the gene body of genes that are two-fold upregulated by exposure to pathogenic PA14. **(E)** Venn diagram of induced genes that are dependent upon *pmk-1* or *atf-7* for complete upregulation.

To evaluate the role of ATF-7 in the direct regulation of genes induced by *P. aeruginosa* infection, we performed chromatin immunoprecipitation followed by sequencing (ChIP-seq) of animals carrying a GFP-tag fused to the C-terminal end of the endogenous *atf-7* locus. Using a GFP polyclonal antibody for immunoprecipitation, we generated ChIP binding profiles for animals in either the wild-type background (*atf-7(qd328[atf-7::2xTY1::GFP])*) or the *pmk-1* mutant background (*pmk-1(km25);atf-7(qd328[atf-7::2xTY1::GFP])*) after a four hour exposure to either *E. coli* OP50 or *P. aeruginosa* PA14, for a total of four conditions, similar to the treatment described in Figure 1A. In all conditions analyzed, ATF-7 exhibited abundant association throughout the genome, with around 9,000 total peaks identified as enriched by MACS2, corresponding to 23% of genes and 25% of transcription start sites (TSSs) (Table S2).

Analysis of the ATF-7 binding profile across all genes associated with enriched TSSs, as well as the subset altered in expression by *P. aeruginosa* in wild-type animals according to our RNA-seq data, revealed that ATF-7 is preferentially located at the promoter regions of genes that are increased in expression by *P. aeruginosa,* and that this enrichment for ATF-7 is lessened by *pmk-1* loss (Figure 2B, Figure S3A). MEME analysis of the most enriched loci identified significant enrichment for the motif GACgTCA, which corresponds to the Jun D bZIP motif expected for ATF-7 (Figure 2A). This motif is present in as many as 80% of the most highly enriched regions of the genome and its abundance is positively correlated with enrichment levels.

**Figure 2:**
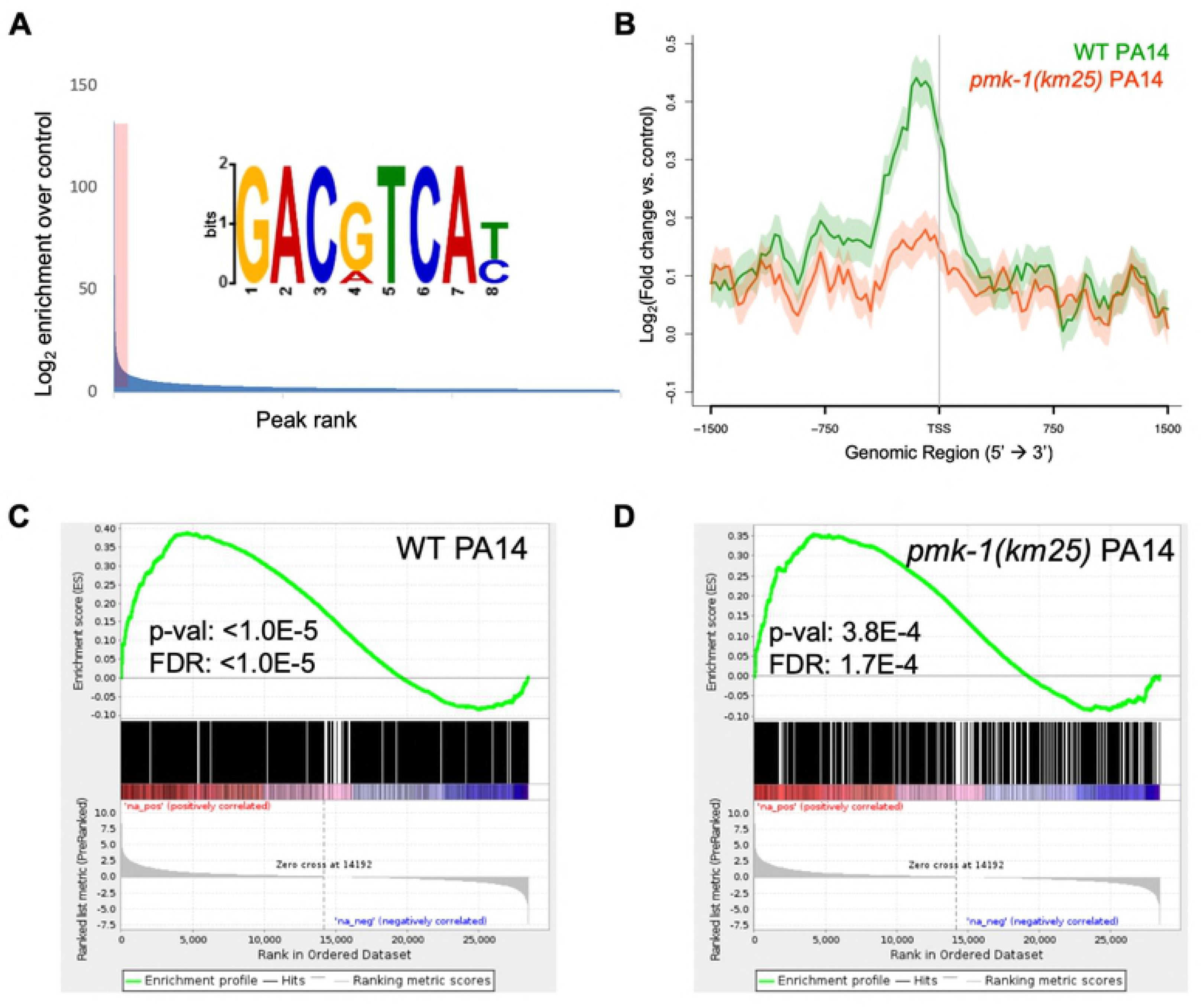
ATF-7 associates with genes that are differentially expressed upon exposure to pathogenic *P. aeruginosa.* **(A)** Motif analysis of ATF-7::GFP ChIP peaks. The top 400 peaks (indicated by red shading) were considered for motif analysis. **(B)** Metagene analysis of ATF-7::GFP binding profile in WT or *pmk-1(km25)* mutant animals exposed to PA14 across genes that are two-fold upregulated (by RNA-seq) upon exposure to PA14. Shading represents standard error among replicates. **(C,D)** Gene Set Enrichment Analysis (GSEA) of transcripts detected by RNA-seq (ranked from most upregulated to most downregulated upon PA14 exposure in N2 animals) for association with ATF-7::GFP peaks in WT (C) or *pmk-1(km25)* mutant (D) animals exposed to PA14.

To identify the most likely immediate downstream targets of ATF-7, we set a peak threshold based on the fraction of peaks containing the bZIP motif after ranking ATF-7 peaks by enrichment (Figure S3B). This resulted in ∼1500-4000 highly enriched locations per experiment. Overlap of the of the retained ATF-7 binding profile of *P. aeruginoasa-*induced genes compared to the RNA-seq data from *P. aeruginosa* induction was measured using a Gene Set Enrichment Analysis (GSEA), which showed that ChIP peaks were enriched for association with transcripts that are positively changed upon pathogen exposure in both *E. coli* and *P. aeruginosa* ChIP conditions (Figure 2C, Figure S3C). This association remains in the *pmk-1* mutant, although at weaker significance level (Figure 2D, Figure S3D). Moreover, we also evaluated ATF-7 binding at individual genomic loci induced by *P. aeruginosa* infection that were dependent on ATF-7 for full upregulation. Examinations of distinct genetic loci further support the conclusions drawn from the metagene analyses described above (Figure 3). These observations suggest a direct transcriptional regulatory role for ATF-7 in the induction of broad transcriptional changes upon immune challenge involving activation of p38/PMK-1 MAPK signaling in response to *P. aeruginosa* infection.

**Figure 3:**
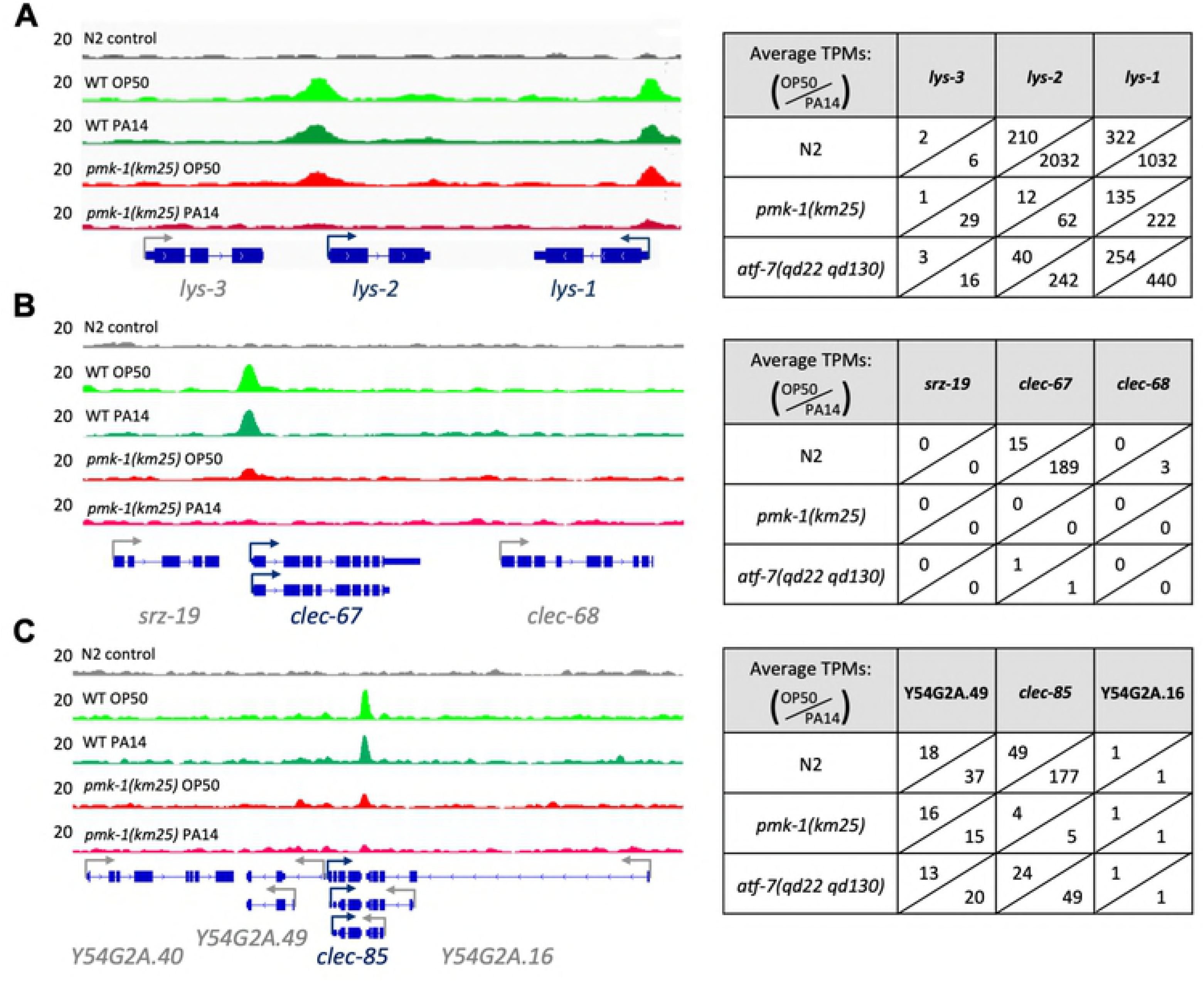
ATF-7 binding at PA14-induced genes. Examples of ATF-7::GFP read pileup at individual loci in all four ChIP conditions. Expression in transcripts per kilobase million (TPM) in all RNA-seq conditions are displayed to the right of each locus. For genes with multiple isoforms, the most highly abundant transcript is displayed.

For functional validation of putative ATF-7-regulated immune response target genes, we focused on transcripts that were upregulated at least two-fold by *P. aeruginosa* exposure in an ATF-7*-*dependent manner and that were also bound by ATF-7 in any of our four ChIP-seq conditions. Included among these putative ATF-7 targets were genes encoding antimicrobial effector molecules, such as CTLD-containing proteins and lysozymes (Table S3). We determined whether RNAi-mediated knockdown of these genes resulted in enhanced susceptibility to killing by *P. aeruginosa* and observed that RNAi of 13 of 43 genes conferred enhanced sensitivity to killing by *P. aeruginosa*, without affecting survival on non-pathogenic *E. coli* (Table S3, Figure S4).

Our data suggest that ATF-7 is a direct regulator of immune effector genes that is regulated by PMK-1 p38 MAPK. We previously proposed a model in which PMK-1 phosphorylates ATF-7 in response to pathogen infection, switching the activity of ATF-7 from that of a transcriptional repressor to that of an activator, allowing the induction of immune response genes [7]. Our data here are consistent with this model, showing a strong dependence of pathogen-induced gene induction on PMK-1 and ATF-7, and a high degree of occupancy of regulatory regions of pathogen-induced genes by ATF-7 under basal and pathogen-induced conditions, with ATF-7 occupancy of pathogen-induced genes being strongly dependent on PMK-1. Moreover, our data reveal that PMK-1-ATF-7 signaling regulates over half of all pathogen-induced genes at the genome-wide level.

PMK-1 signaling has also been implicated in a number of non-infection contexts in *C. elegans* [2,16,17]. Interestingly, we observed that ATF-7 binds quite strongly to several key regulators of stress response pathways. We found that ATF-7 exhibits binding affinity to regulators of autophagy (*lgg-1*), the Unfolded Protein Response (*xbp-1*), and the oxidative stress response (*skn-1*), as well as several immunity regulators (*hlh-30, zip-2*, and interestingly, *atf-7*) (Figure 4). These observations suggest that initiation of other stress responses may be integrated with the immune response. For example, we have previously shown that immune response activation in developing larval is lethal without compensatory XBP-1 activity, establishing an essential role for XBP-1 during activation of innate immunity during infection of *C. elegans* [2]. We speculate that ATF-7 may function to activate anticipatory stress responses that can be activated in concert with innate immunity to promote host survival during microbial infection in a context-dependent manner. Our genomic and genetic findings in the simple, genetically tractable *C. elegans* host reveal a striking degree of global regulation of the organismal response to pathogenic bacteria through a single p38 MAPK-regulated transcriptional regulator. Our data support the idea that host defense, on a genome-wide and organism-wide level, is under the control of a limited number of stress-activated signaling pathways that regulate global regulators of gene transcription.

**Figure 4:**
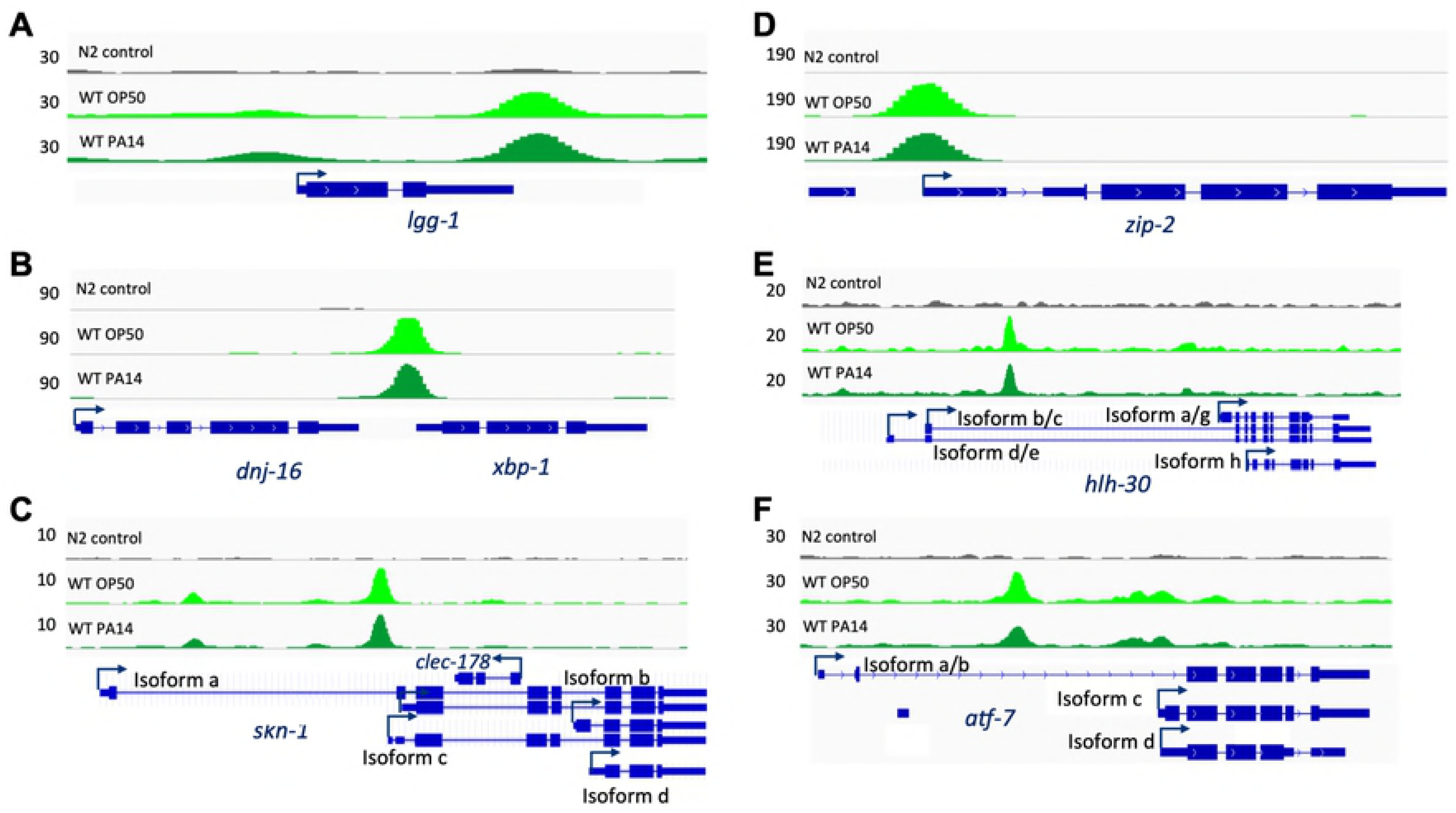
ATF-7 binds to regulatory regions of key regulators of stress physiology. Examples of ATF-7::GFP read pileups stress response (A-C) and immune (D-F) regulators.

## Materials and Methods

### *C. elegans* Strains

Strains used: N2, ZD386 (*atf-7(qd22 qd130)*), KU25 (*pmk-1(km25)*), ZD1807 (*atf-7(qd328[atf-7::2xTY1::GFP])*), ZD1976 (*atf-7(qd328[atf-7::2xTY1::GFP]);pmk-1(km25)*). *C. elegans* were maintained at 16°C on *E. coli* OP50 as described [18]. The *atf-7(qd328)* allele was generated by the CRISPR-Cas9 system as described [19,20] and verified by Sanger sequencing. GFP expression in ZD1807 (*atf-7(qd328)*) was verified by immunobloting, and pull-down was assessed by IP-IB. The *atf-7(qd238)* allele was confirmed to function as wild-type, as assayed by susceptibility to *P. aeruginosa* strain PA14 in a slow kill assay, and then crossed into the *pmk-1(km25)* mutant background.

### Preparation of Animals for Sequencing Experiments

Slow Kill Assay (SKA) plates were prepared as previously described [21]. *P. aeruginosa* strain PA14 or *E. coli* OP50 was grown overnight in Luria Broth (LB), seeded onto SKA media and then grown overnight at 37°C, followed by an additional day at room temperature as previously described [22]. Large populations of animals were synchronized by egg-preparation of gravid adult worms in bleach, followed by L1 arrest overnight in M9 buffer. L1 animals were dropped onto concentrated OP50 lawns seeded onto Nematode Growth Media (NGM) and raised to the L4 larval stage at 20°C (about 40 h). Upon reaching L4, worms were washed off growth plates with M9 and placed on SKA plates prepared as described above, seeded with either PA14 or OP50 and incubated at 25°C for four hours. At this time, worms were harvested by washing for downstream applications.

### Chromatin Immunoprecipitation Followed by Sequencing

After three washes in M9 buffer, animal pellets were resuspended in an equal volume of PBS + complete ULTRA protease inhibitor tablets (Roche), flash frozen in liquid nitrogen, and stored at −80°C until chromatin immunoprecipitation (ChIP). ChIP was preformed as described [23,24] using Ab290, a ChIP-grade polyclonal GFP antibody (Abcam). Libraries were prepared using the SPRIworks Fragment Library System (Beckman Coulter) and single-end sequenced on an Illumina HiSeq2000 sequencer. Three biological replicates of at least 15,000 animals were prepared and sequenced for each condition, with the exception of only two replicates for *atf-7(qd328)* on PA14, as one of the samples failed to pass quality control.

ChIP-seq reads were aligned against the *C. elegans* WBPS9 assembly using bwa v. 0.7.12-r1039 [25] and the resulting bam files were sorted and indexed using samtools v. 1.3 [26]. Sorted bam files were pooled by strain and microbial treatment, and peaks were called using MACS2 (v. 2.1.1.20160309), as follows: callpeak on specific strain bam file (“-t” flag) against the N2_PA14 control sample bam file (“-c” flag) callpeak -c N2_PA14_control.sorted.bam -g ce --keep-dup all --call-summits --extsize 150 -p 1e-3 --nomodel -B. Peak locations were intersected with regions +/-0.5kb around annotated TSS based on the WBPS9/WS258 annotation using bedtools intersect (v2.26.0) [27], and in cases of multiple peaks associated with a given TSS, peaks with maximal enrichment over N2 control were retained. For the purpose of motif identification, peaks were ranked by fold-enrichment over N2 control in descending order and the top 400 peaks were retained, regions +/- 200 bps around the summit were retrieved and sequences were obtained with bedtools getfasta. MEME-ChIP v. 4.12.0 [28] was used to call motifs using the following parameters: meme-chip -oc. -time 300 -order 1 -db db/JASPAR/JASPAR2018_CORE_nematodes_non-redundant.meme -meme-mod anr -meme-minw 5 -meme-maxw 30 -meme-nmotifs 8 -dreme-e 0.05 -centrimo-local -centrimo-score 5.0 -centrimo-ethresh 10.0. Presence of the top motifs under each peak called by macs2 was assessed using Mast v.5.0.1 [29] on the same +/- 200bp region around the summit of each peak. The number of peaks with one or more occurrences of the motif was tallied using a 200-peak window, and plotted across all peaks ranked either by log-fold enrichment over N2 or –log-transformed p-values. Inflection points in the motif density function were used to narrow down the number of peaks retained for downstream analyses.

### RNA Sequencing

After three washes in M9 buffer, TRIzol^TM^ Reagent (Invitrogen) was added to worm pellets and flash frozen in liquid nitrogen. Following an additional round of freeze-thaw, RNA was isolated using the Direct-zol^TM^ RNA MiniPrep kit (Zymo Research). Libraries were prepared using the Kapa mRNA Hyperprep kit and paired end reads were sequenced on the Illumina NextSeq500 sequencer. Three biological replicates of at least 1,000 animals were prepared and sequenced for each condition, with the exception of only two replicates for *atf-7(qd22 qd130)* on PA14, as one of the samples failed to pass quality control.

Reads were aligned against the *C. elegans* WBPS9 assembly/ WS258 annotation using STAR v. 2.5.3a [30] with the following flags: -runThreadN 16 --runMode alignReads --outFilterType BySJout --outFilterMultimapNmax 20 --alignSJoverhangMin 8 --alignSJDBoverhangMin 1 --outFilterMismatchNmax 999 --alignIntronMin 10 --alignIntronMax 1000000 --alignMatesGapMax 1000000 --outSAMtype BAM SortedByCoordinate --quantMode TranscriptomeSAM. with --genomeDir pointing to a low-memory footprint, 75nt-junction WBPS9/WS258 STAR suffix array. Gene expression was quantitated using RSEM v. 1.3.0 [31] with the following flags for all libraries: rsem-calculate-expression --calc-pme --alignments -p 8 against an annotation matching the STAR SA reference. Posterior mean estimates (pme) of counts and estimated “transcript-TPMs” were retrieved for genes and isoforms. Subsequently, counts of isoforms sharing a transcription start site (TSS) were summed, and differential-expression analysis was carried out using DESeq2 [32] in the R v3.4.0 statistical environment, building pairwise models of conditions to be compared (microbial exposures within each genotype). Sequencing library size factors were estimated for each library to account for differences in sequencing depth and complexity among libraries, as well as gene-specific count dispersion parameters (reflecting the relationship between the variance in a given gene’s counts and that gene’s mean expression across samples).

Differences in gene expression between conditions (expressed as log2-transformed fold-changes in expression levels) were estimated under a general linear model (GLM) framework fitted on the read counts. In this model, read counts of each gene in each sample were modeled under a negative binomial distribution, based on the fitted mean of the counts and aforementioned dispersion parameters. Differential expression significance was assessed using a Wald test on the fitted count data (all these steps were performed using the DESeq() function in DESeq2) [32]. P-values were adjusted for multiple-comparison testing using the Benjamini-Hochberg procedure [33].

### Data availability

Raw data presented in this manuscript have been deposited in NCBI’s Gene Expression Omnibus [34] and are accessible through GEO SuperSeries accession number GSE119294, which contains SubSeries GSE119292 (RNA-seq data, including count files) and SubSeries GSE119293 (ChIP-seq data, including wig files and peak calls).

### Evaluation of ATF-7 binding and modulation of gene expression

Metagene analyses of gene expression and ATF-7 binding enrichment were generated by ngs.plot as described [35], using ChIP .bam files from each condition normalized to N2 control as input. Genes considered two-fold upregulated or downregulated are listed in the “N2_up” and “N2_down” tabs of Table S1, respectively.

Correlations between ATF-7 binding and regulation of gene expression were interrogated using the gene set enrichment analysis (GSEA) framework [36]. Briefly, all transcription start sites (TSSs) associated with a protein-coding transcript were ranked based on differential expression results from DESeq2 (log2 fold-changes), which is a measure of the correlation between their expression and the host response to infectious agents. Biases in expression of ATF-7-bound TSSs were assessed using a walk down the list tallying a running-sum statistic, which increases each time a TSS is part of the list and decreases otherwise. The maximum of this metrics (i.e. where the distribution if furthest away from the background) is called the enrichment score (ES). Significance is estimated using random permutations of the TSSs to generate p-values gauging how often the observed ES can be seen in randomized gene sets, for each direction of the expression biases independently. Multiple-testing correction is addressed using a false-discovery rate calculation on permuted datasets.

### Gene Ontology analysis

Genes with adjusted p-values <0.05 were considered for Gene Ontology enrichment analysis using the DAVID online webtool, considering as a background the union of all genes with a non-zero baseMean value across any of the DE comparison, based on unique WormBase IDs.

### Killing Assays and Bacterial Strains

PA14 plates were prepared as described as above. N2 animals were grown on NGM, supplemented with 25 ug/mL carbenicillin and 2mM isopropyl b-D-1 thiogalactopyranoside (IPTG), that was seeded with either the *E. coli* HT115 expressing plasmids targeting the gene of interest or the empty L4440 vector backbone for two generations prior to each experiment. Animal populations were synchronized by egg lay. At the L4 larval stage, approximately 30 worms were transferred to prepared SKA plates and incubated at 25°C. Animals were scored for killing twice daily until the majority of animals had died. Within each experiment, three plates were prepared and scored per RNAi treatment. All clones were obtained from the Ahringer [37] or Vidal [38]RNAi libraries and were verified by sequencing. For a list of all RNAi clones used, see Table S4.

## Author Contributions

M.F. and E.J.T. performed all experiments. V.B. and S.S.L. performed bioinformatics analysis of RNA-seq and ChIP-seq datasets. M.F. and D.H.K analyzed data, interpreted results, and wrote the paper with input from E.J.T.

## Acknowledgements

We thank H.R. Horvitz and the *Caenorhabditis* Genetics Center for providing strains and reagents. This work was funded by NIH grants R01GM084477 (to DHK) and T32GM007287 (to MF and EJT).

---

**Supplemental Figure 1:**
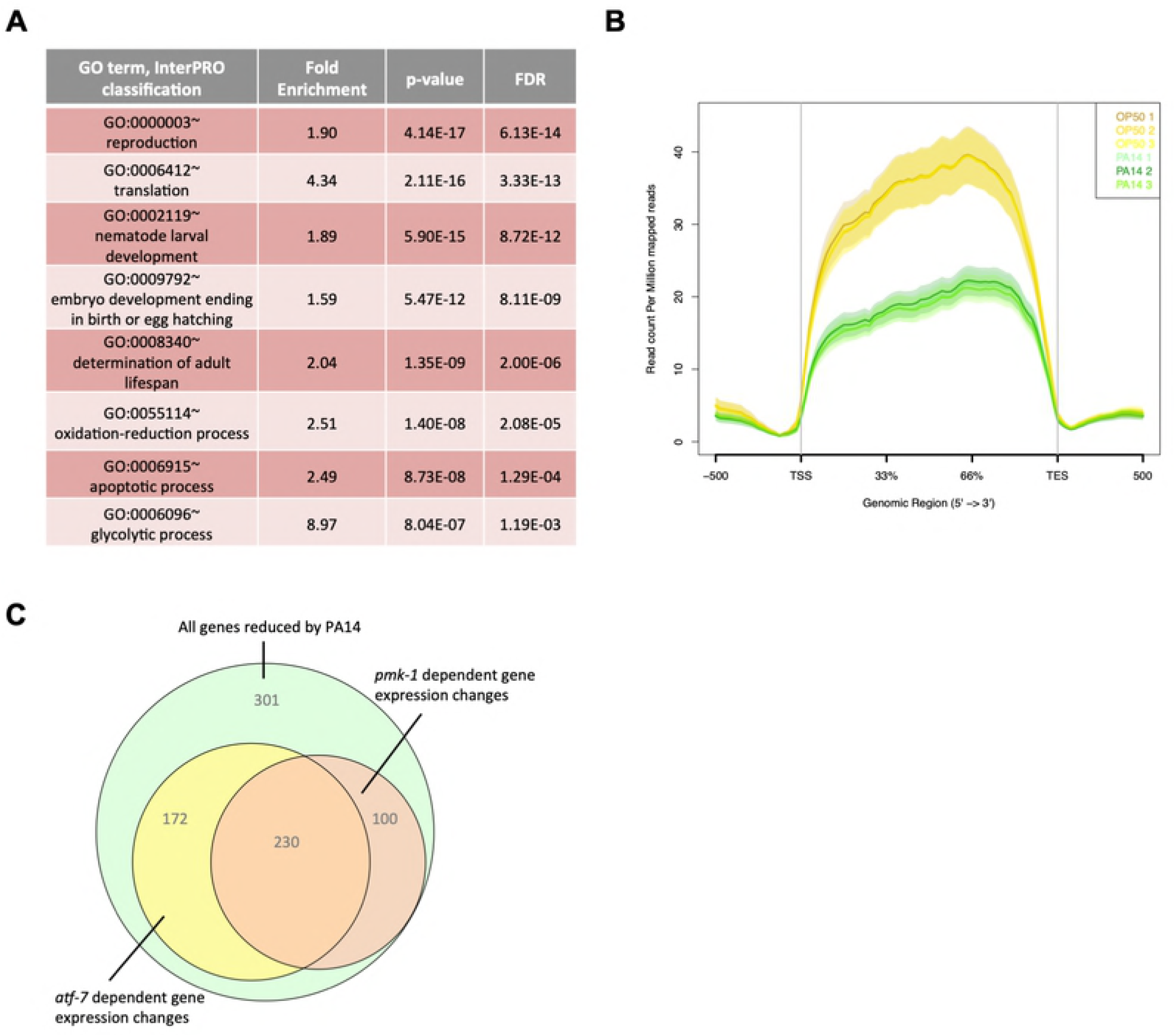
Expression of genes decreased by PA14 exposure. **(A)** Top GO terms and InterPRO classifications of transcripts that are significantly downregulated (adjusted p-value < 0.05) in N2 animals exposed to PA14 versus OP50. **(B)** Average expression (RPM) across the gene body of genes that are two-fold downregulated by exposure to pathogenic PA14. **(C)** Venn diagram of decreased genes that are dependent upon *pmk-1* or *atf-7* for complete downregulation.

**Supplemental Figure 2:**
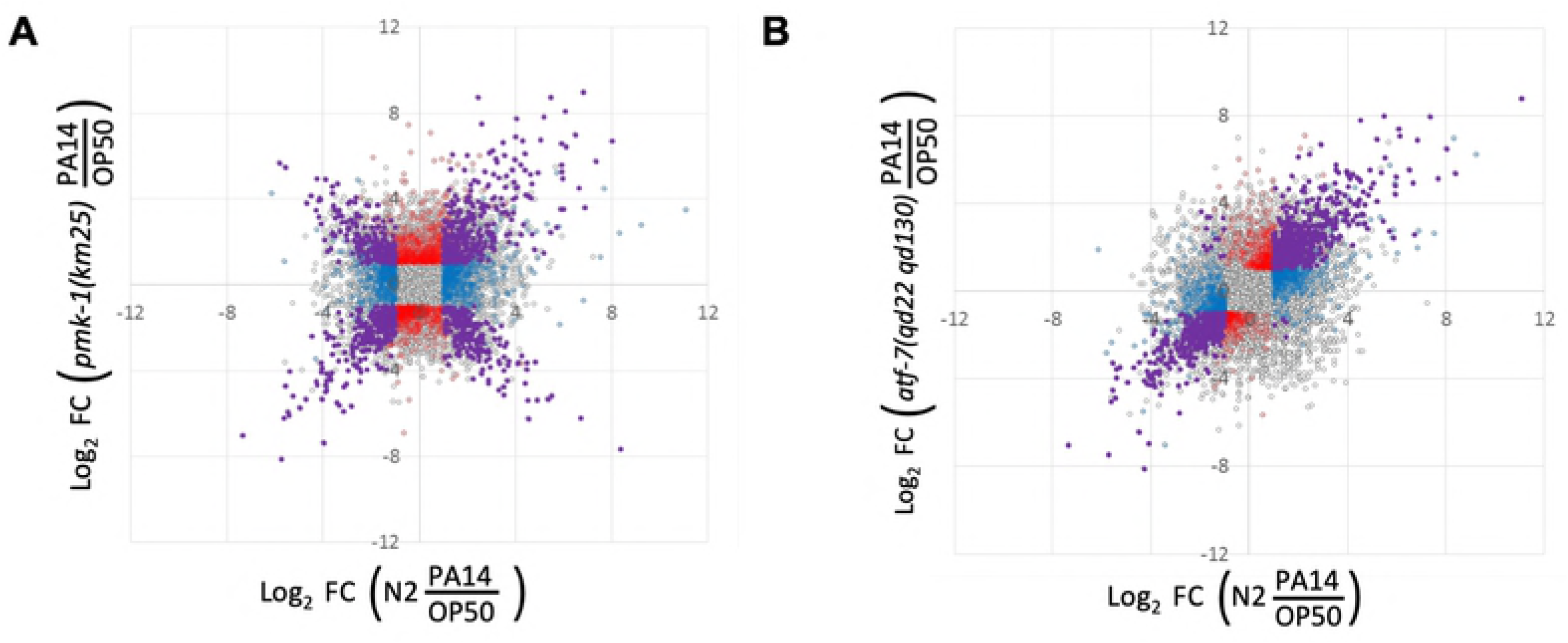
Differential PA14-induced gene expression in *pmk-1* and *atf-7* mutant animals. 2×2 comparison of genes differentially expressed by exposure to PA14 in N2 animals (x-axis) or upon loss of *pmk-1* (A) or *atf-7* (B) (y-axis). Transcripts highlighted in purple correspond to genes that were significantly different compared to OP50 in both genotypes (adjusted p-value of <0.05). Blue dots indicate genes that are significantly different in the N2 PA14/OP50 comparison only. Red dots represent genes that reach significance in only the mutant condition. Grey dots indicate genes with detected transcripts in at least one condition being compared, but that failed to reach significance cutoffs in either data set.

**Supplemental Figure 3:**
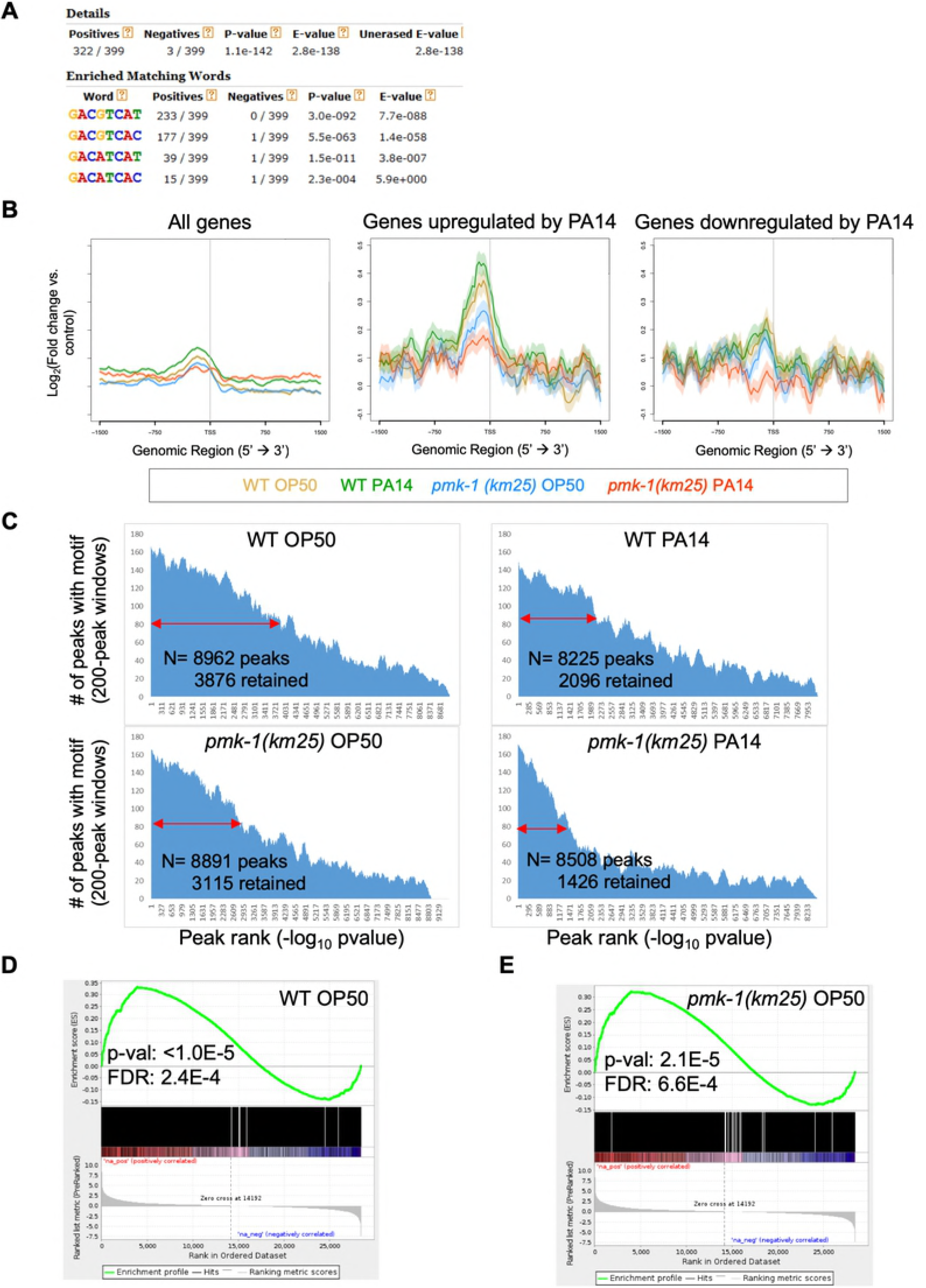
Evaluation of ATF-7::GFP peaks. **(A)** Motif analysis of ATF-7::GFP ChIP peaks. The top 400 peaks were considered for motif analysis. **(B)** Metagene analysis of ATF-7::GFP binding in all four ChIP conditions across all genes, genes that are two-fold upregulated, or two-fold downregulated by RNA-seq upon exposure to PA14 in a wild-type (N2) background. Shading represents standard error among replicates. **(C)** Ranked ATF-7::GFP peaks called in animals in all four ChIP conditions. Double red arrow indicates the peaks that were retained for further analysis. **(D,E)** Gene Set Enrichment Analysis (GSEA) of transcripts detected by RNAseq (ranked from most upregulated to most downregulated upon PA14 exposure in N2 animals) for association with ATF-7::GFP peaks in WT (C) or *pmk-1(km25)* mutant (D) animals exposed to OP50.

**Supplemental Figure 4:**
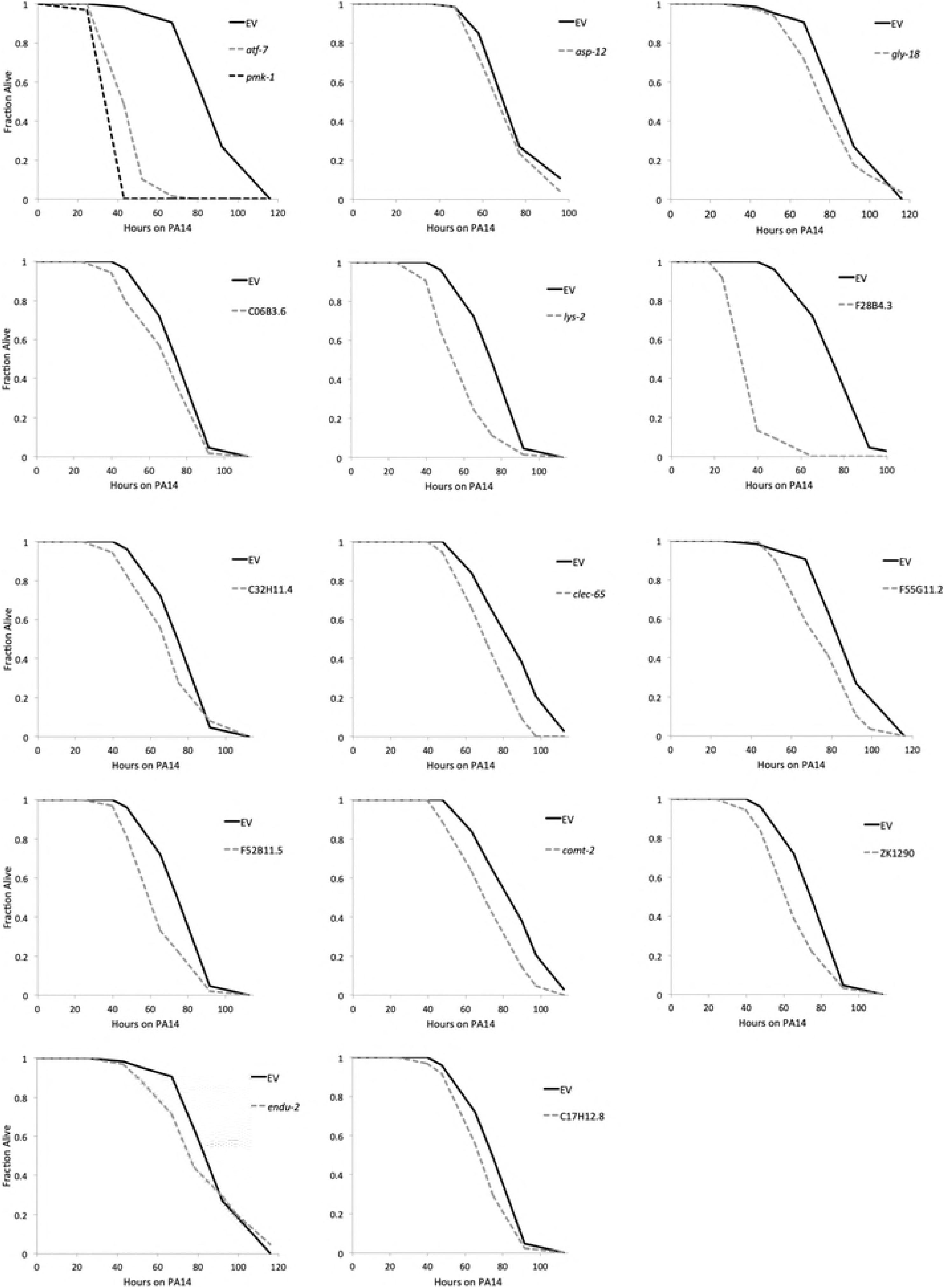
ATF-7 target genes that effect survival on *P. aeruginosa* PA14. Representative survival curves of animals treated with RNAi against indicated genes that resulted in a significant (p-value < 0.05 by log-rank test) reduction in survival on PA14 compared to EV controls in 2/2 experiments. Animals were treated with RNAi for two generations prior to exposure to PA14. EV refers to HT115 carrying the Empty Vector control plasmid, L4440.

**Table S1: RNA-seq summary**

See separate electronic (.xlsx) file.

**Table S2: ATF-7::GFP peaks from ChIP-seq.**

See separate electronic (.xlsx) file.

**Table S3: Genes tested for Esp phenotype by RNAi knockdown.**

Protein domains classified using the David 6.8 Functional Annotation Tool. “Yes,” indicates a significant (p-value < 0.05 by log-rank test) reduction in survival on PA14 compared to Empty Vector control in 2/2 experiments.

